# Expectation of the intercept from bivariate LD score regression in the presence of population stratification

**DOI:** 10.1101/310565

**Authors:** Loic Yengo, Jian Yang, Peter M. Visscher

## Abstract

Linkage disequilibrium (LD) score regression is an increasingly popular method used to quantify the level of confounding in genome-wide association studies (GWAS) or to estimate heritability and genetic correlation between traits. When applied to a pair of GWAS, the LD score regression (LDSC) methodology produces a statistic, referred to as the bivariate LDSC intercept, which deviation from 0 is classically interpreted as an indication of sample overlap between the two GWAS. Here we propose an extension of the theory underlying the bivariate LDSC methodology, which accounts for population stratification within and between GWAS. Our extended theory predicts an inflation of the bivariate LDSC intercept when sample sizes and heritability are large, even in the absence of sample overlap. We illustrate our theoretical results with simulations based on actual SNP genotypes and we propose a re-interpretation of previously published results in the light of our extended theory.

---

Initially introduced in Bulik-Sullivan *et al.* (2014)^1^, the LD score regression (LDSC) methodology relies on a derivation for a particular SNP *j* of the expectation of its association χ^2^-statistic 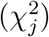 as function of the LD score (*ℓ_j_*), that is the sum of pairwise squared correlation between minor allele counts at SNP *j* versus all SNPs or versus SNPs in its vicinity, the heritability (*h*^2^) of the trait plus a term indicating the level of confounding in the GWAS attributable to population stratification. The derivation proposed in Bulik-Sullivan *et al.* (2014) predicts that the latter term (a.k.a the LDSC intercept) increases linearly with the sample size (*N*) of the GWAS and the heritability of the trait, which raises a number of challenges in its interpretation as an indication of confounding when *N* is very large (*N* ~400,000 for example). This problematic behavior of the LDSC intercept has been underlined in a recent publication by Loh *et al.* (2017)^2^, which recommends the use of an alternative statistic to quantify the influence of population stratification on GWAS results.

The LDSC methodology was later extended in Bulik-Sullivan *et al.* (2015)^3^ to analyse pairs of GWAS in order to quantify the genetic correlation between focus traits of each GWAS. The theory proposed in Bulik-Sullivan *et al.* (2015) introduced the bivariate LD score intercept, obtained from the regression of the product of association statistics (defined further below) onto LD scores, as a measure of sample overlap between the two GWAS. Compared to Bulik-Sullivan *et al.* (2014), the model underlying the bivariate LDSC methodology does not consider the presence of population stratification within and between studies, neither discusses how this might contribute to the expectation of the bivariate LDSC intercept. We therefore address this question here and propose an extension of the initial theory.

## Theoretical results

### Notations

We consider that GWASs of two traits y_1_ and y_2_ are performed in two cohorts: cohort 1 and cohort 2 respectively. Each SNP *j* is tested for association with each trait using a test statistic *T_j_* defined as the ratio between estimated SNP effect from linear regression of y_1_ or y_2_ onto the minor allele count (MAC) over its estimated standard error. Let us denote *T*_*j*1_ and *T*_*j*2_ the test statistics for SNP *j* calculated for y_1_ in cohort 1 and for y_2_ in cohort 2 respectively. We propose hereafter an extension of the expectation of the product of *T*_*j*1_ and *T*_*j*2_, initially proposed in Bulik-Sullivan *et al.* (2015), in the presence of population stratification within each cohort induced by genetic drift.

As Bulik-Sullivan et al. (2014), we assume that population stratification can be modelled in each cohort as a 50:50 mixture of two sub-populations deriving from a common ancestral population and thus having different allele frequencies spectra. We denote *σ*_*S*1_ and *σ*_*S*2_ as the mean phenotypic difference between sub-populations of cohort 1 and 2 respectively (environmental stratification). We introduce 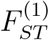 and 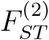 as Wright’s *F_ST_* measures of allele frequency differences between sub-populations of cohort 1 and cohort 2 respectively (genetic stratification). For the sake of simplicity, we assume that the level of genetic and environmental stratification is similar between cohort 1 and cohort 2. Therefore 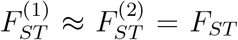 and *σ*_*S*1_ ≈ *σ*_*S*2_ = *σ*_S_. We finally denote 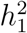, 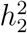 and *r_g_*, the heritabilities of y_1_ and y_2_ and their genetic correlation.

We consider the following model

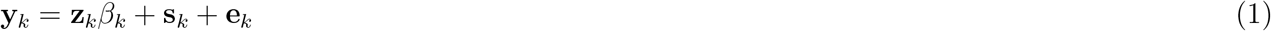

where y_*k*_ is the vector of phenotypes of *N_k_* individuals from cohort *k*, *s_k_* a *N_k_*-dimensional vector which entries equal ±*σ_S_* (mean of environmental fixed effects) depending on whether participants enrolled in cohort *k* are from one or the other sub-population, z_*k*_ a *N_k_* × *M* matrix of scaled MAC, *β*_*k*_ a *M* dimensional vector of true SNP effect sizes on trait y_*k*_ and e_*k*_ a *N_k_*-dimensional vector of residuals.

We denote 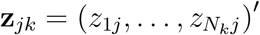 the *j*-th column of matrix z_*k*_. The *i*-th entry of z_*jk*_, denoted *z_ijk_*, is defined as 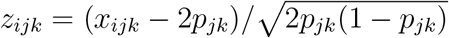 where *x_ijk_* is the MAC at SNP *j* of individual *i* from cohort *k* and *p_jk_* is minor allele frequency (MAF) of SNP *j* in cohort *k*. We assume that all sub-samples of cohorts 1 and 2 have derived from the same ancestral population and denote *p_j_* as the MAF of SNP *j* in that ancestral population.

As in Bulik-Sullivan *et al.* (2015), we model the true genetic effects as

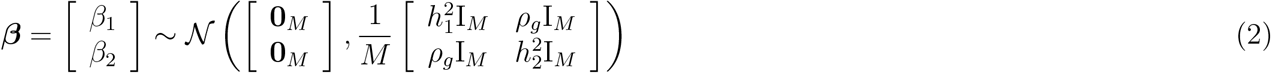

and the residuals as

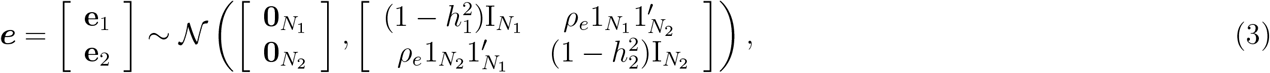

where 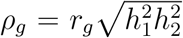 is the genetic covariance between y_1_ and y_2_ and *ρ_e_* is the covariance between error terms e_1_ and e_2_. We also define *ρ* as *ρ* ≡ *ρ_g_* + *ρ_e_*.

In cohort *k*, the least-squares estimate of the effect of SNP *j* on y_*k*_, denoted 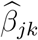, is approximately equal to 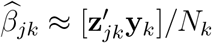 and has a sampling variance ≈ 1/*N_k_*. The accuracy of these approximations increases with the sample size *N_k_*, that we assume here to be large, e.g. hundreds of thousands. Therefore the *t*-statistic *T_jk_* can be approximated as 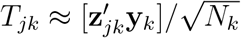.

### Derivation of 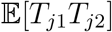

We can express 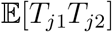 as follows

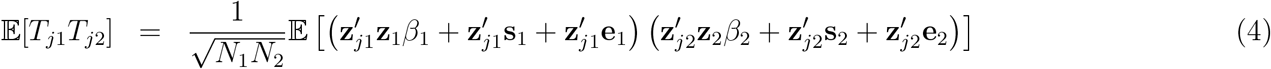

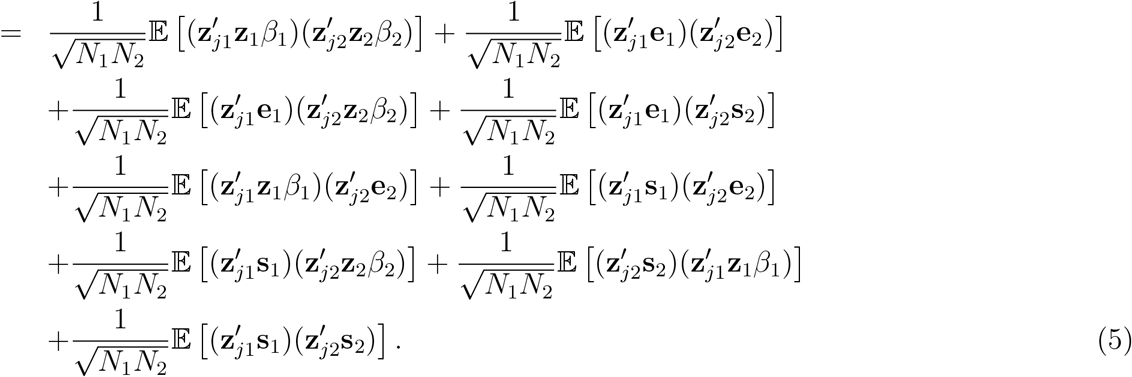

If we assume independence between e_*k*_ and *β_k_*, between e_*k*_ and s_*k*_, and that 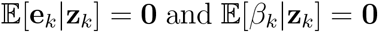, which are classical assumptions made in Bulik-Sullivan *et al*. (2014), then

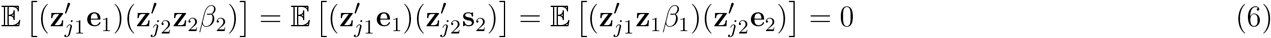

and

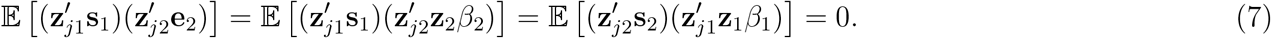

This therefore leads to simplify 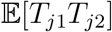 as

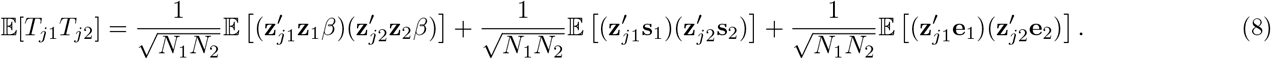

Equation (8) shows a strong similarity with equation (1) from Bulik-Sullivan *et al.* (2015) but one can already notice the inclusion of the second term on the right side of the equation, representing the contribution of population stratification within and between cohorts. We now further simplify equation (8).

Let us start with the first term of the right side of equation (8). We can rewrite it as

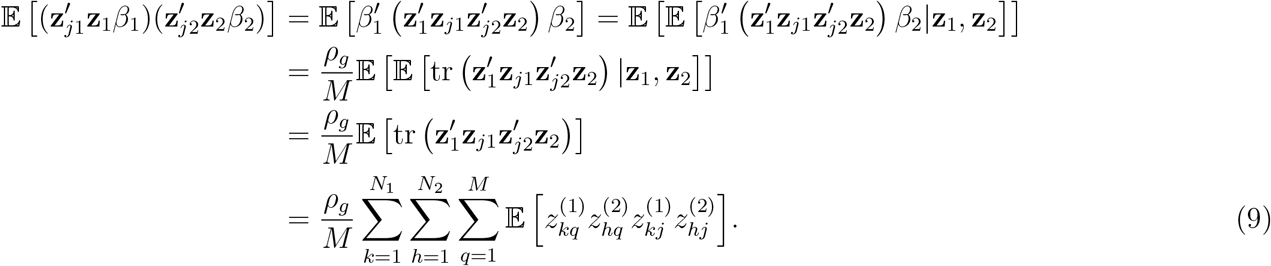

Denote 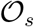 as the set of samples overlapping cohort 1 and cohort 2. We use the simplified notation “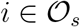” (or “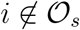”) to indicate that individual *i* belongs (or does not belong) to 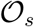. We can therefore write

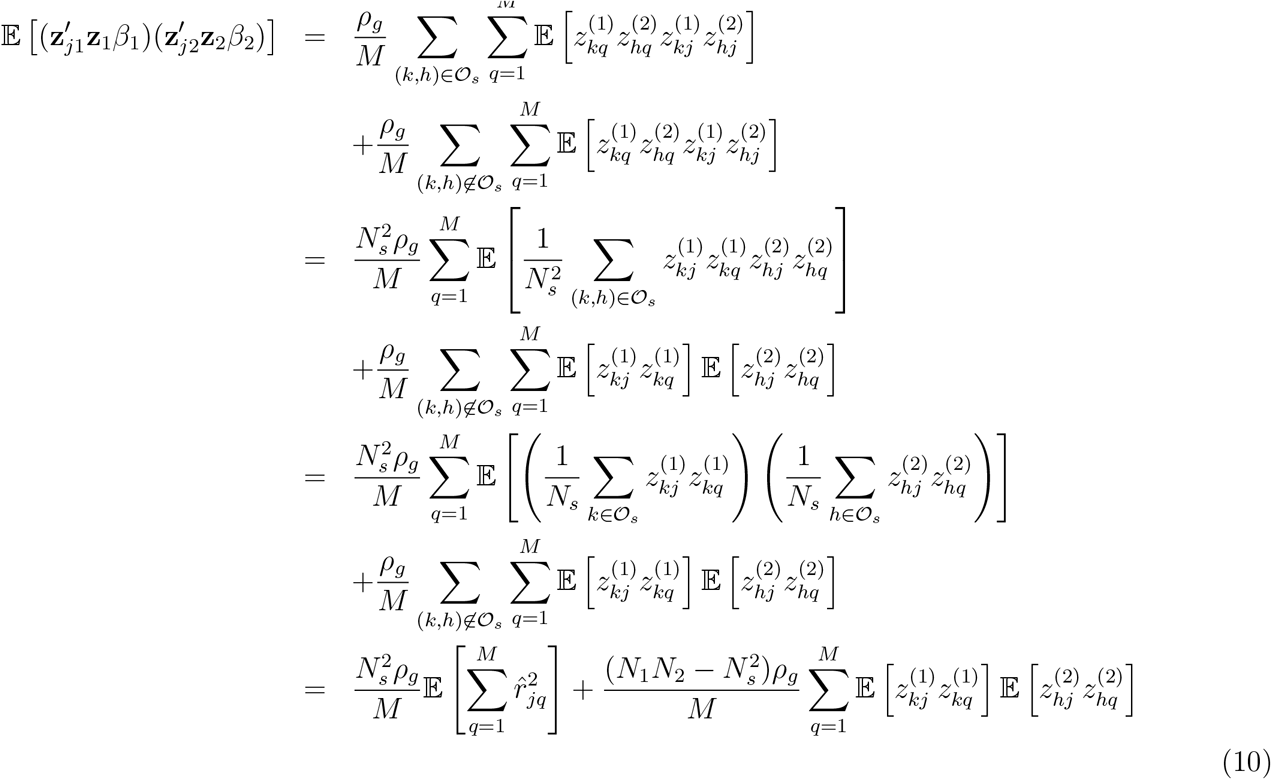

where 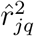 is the squared sample correlation between MAC at SNP *j* and MAC at SNP *q*.

Besides, 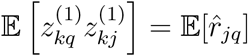 is the expectation of the sample correlation in cohort 1 between MAC at SNP *j* and MAC at SNP *q*, which we assumed to be the same as the expectation of the sample correlation in cohort 2, i.e. 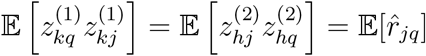.

Therefore 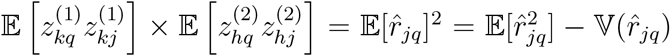. From equation (2.3) in Bulik-Sullivan *et al*. (2014) we can therefore deduce that

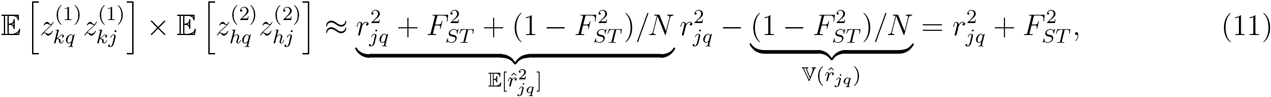

where, in the equation above, *N* = *N*_1_ or *N* = *N*_2_ indifferently. We can therefore rewrite equation (10) as

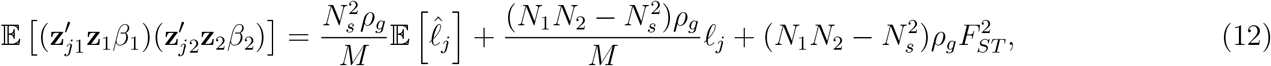

where 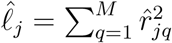 is the sample LD score of SNP *j* calculated only from samples in 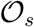 and 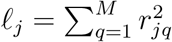 is theoretical true LD score.

If we assume that 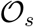 is a random sample of cohort 1 and cohort 2, then the population structure within 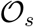 is expected to be similar to that within cohort 1 and within cohort 2. Bulik-Sullivan *et al.* (2014) derived an approximation of the expectation of the sample LD score 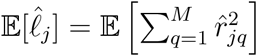 (equation 2.4 of their supplementary note) that we rewrite here as

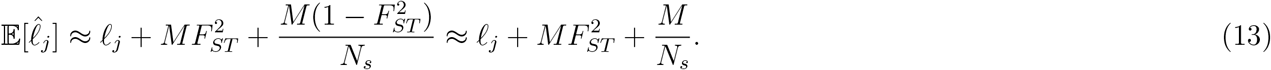

If we combine equations (12) and (13) we get

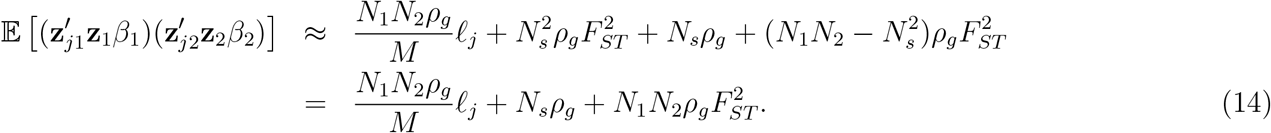

From Bulik-Sullivan *et al.* (2015) we know that 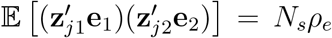. Therefore from combining equations (8) and (14) we have

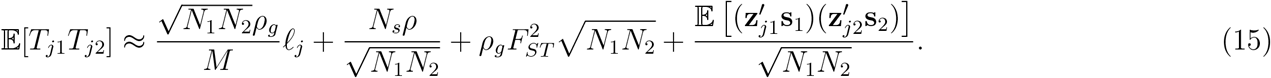

The last term to derive, 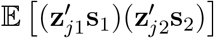, can be rewritten as in Bulik-Sullivan *et al.* (2014) (eq. 2.11) as

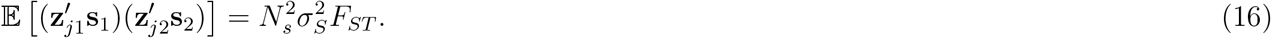

By combining equations (15) and (16), we obtain the final expression

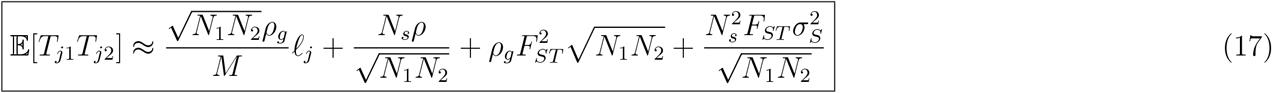

As a special case we can obtain the univariate LD score regression equation by assuming that cohort 1 is the same as cohort 2, i.e. *N*_1_ = *N*_2_ = *N_s_* = *N* and *ρ* = 1. We hence have

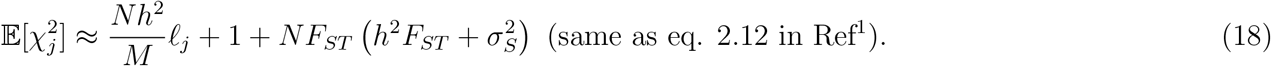

Another interesting special case is the absence of sample overlap (*N_s_* = 0), which leads to

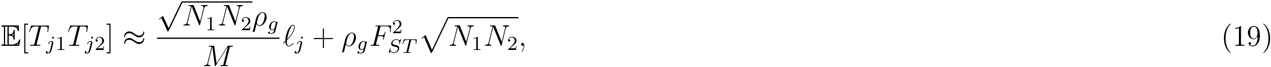

i.e. a non-zero intercept is expected even in the absence of sample overlap. Note in this case that the intercept is proportional to the geometric mean of the sample sizes of both GWAS 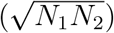 and the genetic covariance between the traits. It is therefore expected to increase with large sample sizes and more heritable traits. As a numerical example, although 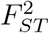 is small in general, values of 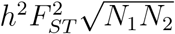 can be as large as ~0.17, which would be indicative of sample overlap if we take for example *h*^2^ = 0.5, *F_ST_* = 0.001, *N*_1_ = 450, 000 (sample size of UK Biobank) and *N*_2_ = 250, 000 (sample size of Wood *et al.*, 2014).

## Simulations

We performed a simulation to quantify the inflation of the bivariate LDSC intercept created when the sample size of each GWAS and the heritability are large. We used for our simulations genotypes at 1,123,348 HapMap 3 SNPs (Online methods) from 348,502 unrelated (genetic relationship < 0.05) participants of the UK Biobank (UKB) with European ancestry (Online methods). To mimic independent GWAS, we randomly split our dataset in two sub-samples of equal size (*N*_1_ = *N*_2_ = 174, 251), and simulated traits from 10,000 causal variants (randomly sampled among HapMap 3 SNPs) and with an heritability varying from 0.1, 0.2,…, up to 0.9. Each trait was simulated with same SNPs effect sizes in each sub-sample so that the genetic correlation is expected to be *r_g_* = 1. For each simulation replicate, we performed a GWAS of each simulated trait in each sub-sample separately, then used GWAS summary statistics to perform a bivariate LD score regression. LD score regression was performed using the LDSC software v1.0.0 and using LD scores from European samples of the 1,000 genomes reference panel. We performed 100 simulation replicates for each value of expected heritability.

We present the results our this simulation in Figure 1. Overall, we found an inflation of the bivariate LDSC intercept, which increases with the heritability of the trait. For example, with *h*^2^ = 0.9 we observed bivariate LDSC intercepts as large as ~0.1 (s.e. 0.02), which under the theory developed in Bulik-Sullivan *et al.* (2015), would falsely indicate a potential overlap of 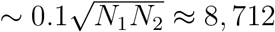 participants between the two sub-samples of the UKB. We also observed an inflation of the univariate LDSC intercept (Figure 1, panel **b**), which is expected under Bulik-Sullivan *et al.* (2014) theory.

**Figure 1:**
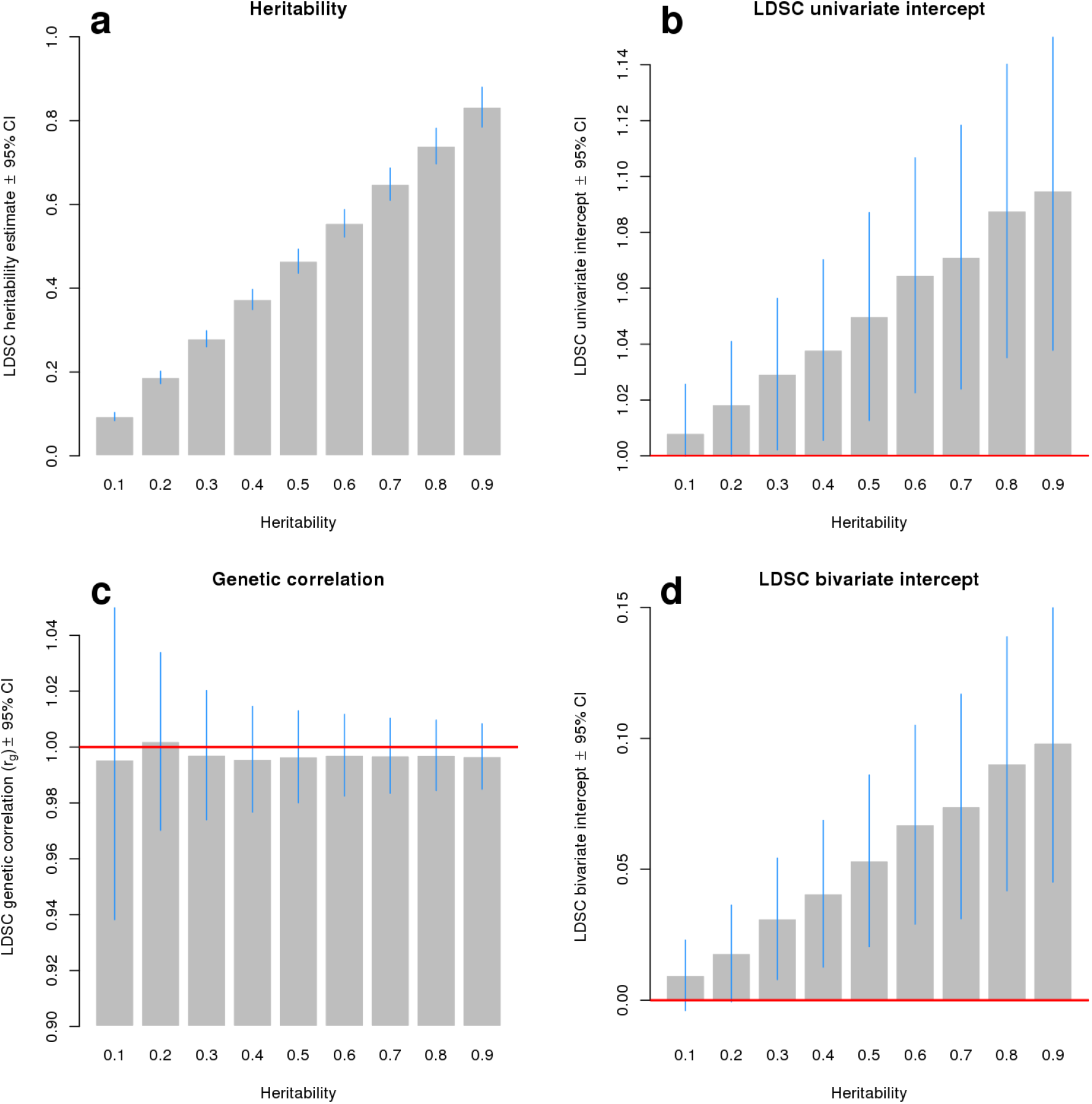
Statistics from the LD score regression applied to 900 simulated GWAS. Panels **a** and **b** show univariate LD score regression intercepts and estimates of heritability respectively obtained from analyzing summary statistics from each sub-sample separately, then averaged between the two independent sub-samples of participants of the UK Biobank (UKB). Panels **c** and **d** show estimates of genetic correlations (expected to be *r_g_* = 1) between the two sub-samples and bivariate LD score regression intercepts respectively, indicating sample overlap between the two sub-samples of UKB, in particular when the underlying heritability is > 0.5.

Under the assumptions made in this simulation 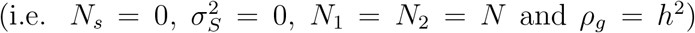, Equation (17) predicts a affine relationship between univariate LDSC intercepts (*I_u_*) within each cohort and bivariate LDSC intercept (*I_b_*): *I_u_* – 1. We validated this prediction in our simulated data as shown on Figure 2.

**Figure 2:**
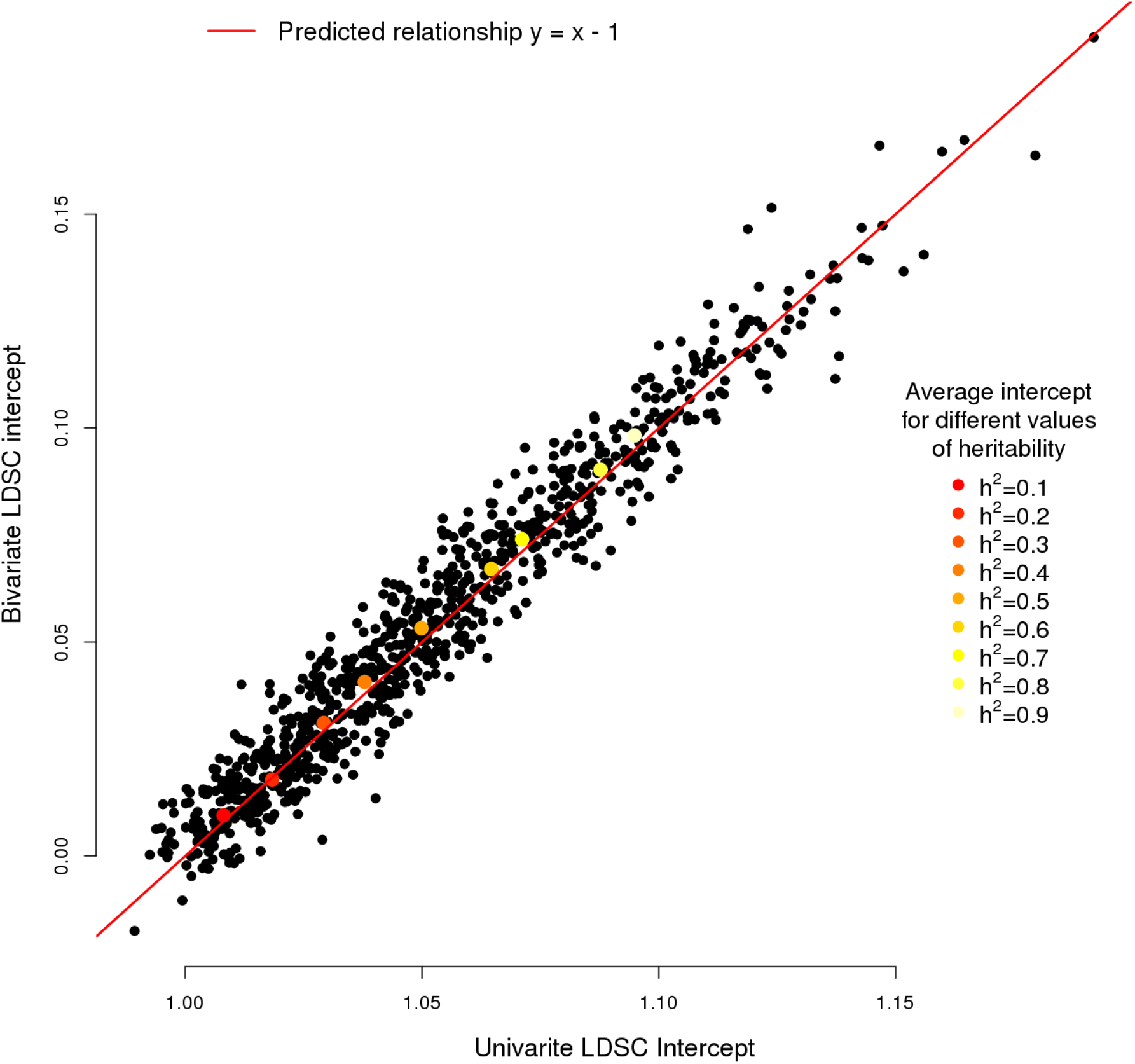
Relationship between univariate (x-axis) and bivariate (y-axis) LD score regression intercepts under the assumption that the same trait is analysed in both cohort (i.e. the genetic correlation *r_g_* = 1), that population stratification does not explain any phenotypic variance 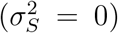 and in the absence of sample overlap (*N_s_* = 0). Each black dot corresponds one simulation replicate as described in our simulation study. Colored dots represent the mean of 100 simulations replicates obtained with a fixed value of heritability (*h*^2^).

## Empirical results: GWAS of height and body mass index (BMI)

We used summary statistics from published GWAS of height^4^ (Wood *et al.*, 2014; median *N* ≈ 252, 083) and BMI^5^ (Locke *et al.*, 2015; median N ¾ 233,692). We also performed a GWAS of height and BMI in 348,502 unrelated participants of the UK Biobank (UKB) of European ancestry. We then estimated the bivariate LDSC intercept obtained from the comparison of Wood *et al.* (2014) with GWAS of height in UKB as well as from the comparison of Locke *et al.* (2015) with GWAS of BMI in UKB. We found in the first case a bivariate LDSC intercept ~0.15 (s.e. 0.04) for height and ~0.01 (0.01) for BMI. We previously reported similar observations using test statistics from linear mixed model association analyses in the UKB (Yengo *et al.*, 2018)^6^. Under Bulik-Sullivan *et al.* (2015) theory, these estimates suggest a significant overlap of 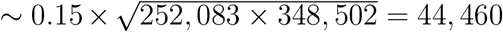 participants between the Wood *et al.* (2014) study and UKB but no signigicant overlap between partcipants of the Locke *et al.* (2015) study and UKB. Given that the same cohorts are included in the Locke *et al.* (2015) study and in the Wood *et al.* (2014) GWAS, these two conclusions are therefore contradictory or inconsistent with the Bulik-Sullivan *et al.* (2015) theory.

To further illustrate this contradiction, we performed in the 348,502 unrelated UKB participants a GWAS of height in females (*N*_1_ = 188, 465) and males (*N*_2_ = 160, 037) separately. Although no true sample overlap is to be expected, we nonetheless found a significant bivariate LDSC intercept of ~ 0.1 (s.e. 0.02), which also suggests a significant sample overlap under Bulik-Sullivan *et al.* (2015) theory.

In the light of the theoretical extension proposed in this note, we believe that these observations can be explained by the large sample sizes considered here, by the difference of heritability between height and BMI and the amount of trait variance explained by population stratification.

## Discussion

We have developed in this note an extension of the theory underlying the bivariate LD score regression methodology in the presence of population stratification within each GWAS. Beyond the Bulik-Sullivan *et al.* (2015) theory, our results show that a non-zero bivariate LDSC intercept does not always indicate sample overlap but may also reflect patterns of population stratification within each study that are shared between studies.

Our extended theory thus explains and predicts a series of puzzling observations that the initial theory does not. For example, we can explain inconsistent detection of sample overlap from GWAS of traits with different heritabilities. Other theoretical extensions of the LDSC methodology have been previously proposed. We may for instance refer to the works of Lu *et al.* (2017)^7^ (GNOVA method) who generalized the ‘‘sample overlap” term (*N_s_*) by replacing it with the sum of genetic relatedness coefficients between participants of the two studies. This extension therefore predicts an inflation of the bivariate LDSC intercept if relatives span both studies, which is more general than the restriction to actual sample overlap. Another contribution by Lee *et al.* (2018)^8^ is also worth mentioning here as it provides a rigorous mathematical framework that not only refines our understanding of the LD score regression methodology but also helps clarifying its interpretation. These two examples, among many others, both illustrate the effervescence of researches driven by the LD score regression methodology.

In conclusion, our findings improve the interpretation of results from bivariate LDSC analyses and may further have implications in other methodologies which use output statistics from LD score regression in their inference.

## Online methods

### UK Biobank data and summary statistics

We used genotypic and phenotypic data (height and body mass index) measured in participants of the UK Biobank. Samples and SNPs selection have been described in a previous publication^6^. We also analysed publicly available GWAS summary statistics from the Wood *et al.* (2014) and Locke *et al.* (2015) GWAS of height and BM respectively. Summary statistics from these studies were downloaded from the following link: https://portals.broadinstitute.org/collaboration/giant/index.php/GIANT_consortium_data_files.

### GWAS simulation

We simulated 900 GWAS (9 values of heritabiliy × 100 simulation replicates) according to the following steps: 1) For each simulation replicate and each value of heritability, we randomly sampled *M* = 10, 000 SNPs as causal variants. 2) For each of the 348,502 UKB unrelated participants, we then simulated a quantitive trait y using the following equation

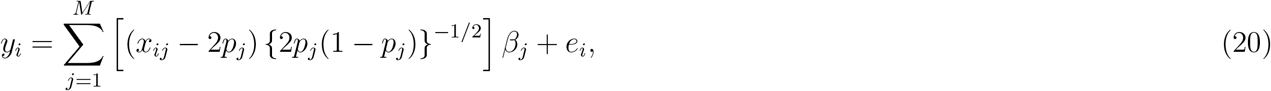

where *X_j_* is the MAC of individual *i* at SNP *j*, *p_j_* is the MAF of SNP *j* and *β_j_* and *e_i_* are independent normally distributed terms such as

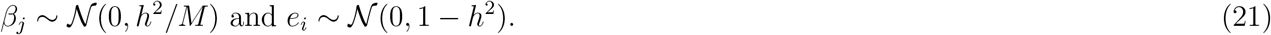

3) Once the trait is simulated we randomly split the cohort into 2 equally sized sub-cohorts and performed SNP-trait association analyses using PLINK^9^ in each sub-cohort.

## Acknowledgements

This research was supported by the Australian Research Council (DP130102666 and DP160103860), the Australian National Health and Medical Research Council (1078037, 1078901, 1103418, 1107258, 1127440 and 1113400). This research has been conducted using the UK Biobank Resource under project 12505. We thank Tim Frayling, Joel Hirschhorn and members of GIANT consortium for fruitful discussions.

